# Uncertain Precision: Neurobiological and Physiological Correlates of Conservative Confidence Bias

**DOI:** 10.1101/2022.08.01.502365

**Authors:** Micah Allen, Tobias U. Hauser, Dietrich Samuel Schwarzkopf, Raymond J. Dolan, Geraint Rees

## Abstract

Correctly estimating the influence of uncertainty on our decisions is a critical metacognitive faculty. However, the relationship between sensory uncertainty (or its inverse, precision), decision accuracy, and subjective confidence is currently unclear. Although some findings indicate that healthy adults exhibit an illusion of over-confidence, under-confidence in response to sensory uncertainty has also been reported. One reason for this ambiguity is that stimulus intensity and precision are typically confounded with one another, limiting the ability to assess their independent contribution to metacognitive biases. Here we report four psychophysical experiments controlling these factors, finding that healthy human participants are systematically under-confident when discriminating low-precision stimuli. This bias remains even when decision accuracy and reaction time are accounted for, indicating that a performance-independent computation partially underpins the influence of sensory precision on confidence. We further show that this influence is linked to fluctuations in arousal and individual differences in the neuroanatomy of the left superior parietal lobe and middle insula. These results illuminate the neural and physiological correlates of precision misperception in metacognition.

**Significance Statement:** The ability to recognize the influence of sensory uncertainty on our decisions underpins the veracity of self-monitoring, or metacognition. In the extreme, a systematic confidence bias can undermine decision accuracy and potentially underpin disordered self-insight in neuropsychiatric illness. Previously it was unclear if metacognition accurately reflects changes in sensory precision, in part due to confounding effects of stimulus intensity and precision. Here we overcome these limitations to repeatedly demonstrate a robust precision-related confidence bias. Further, we reveal novel neuroanatomical and physiological markers underlying this metacognitive bias. These results suggest a unique state-based computational mechanism may drive subjective confidence biases and further provide new avenues for investigating maladaptive awareness of uncertainty in neuropsychiatric disorders.

## Introduction

Our perception of the external world is inherently uncertain. This uncertainty stems both from the volatility of the world itself, and from the noisy biomechanical processes by which sensory inputs are transcribed into neural signals. For example, my perception of a lighthouse on the horizon can be mistaken either because my sensory information is uncertain (e.g., my vision is obscured by fog), or because I myself have become an unreliable observer - perhaps I am tired, and my eyesight has weakened. In either case, if I do not recognize the influence of uncertainty on my perception, then I am unlikely to learn from my mistakes and will make increasingly poor decisions, colliding with unseen calamity. In this sense, the metacognitive ability to monitor sensory uncertainty (or it’s inverse, precision) is a crucial aspect of perceptual decision-making.

Currently there is substantial ambiguity as to how, and even if, healthy human participants embed sensory precision in their decision confidence. In the lab, psychophysical random-dot motion stimuli provide an ideal opportunity to manipulate sensory precision (i.e., the distribution of dot motions) independently from signal intensity (i.e., the overall mean direction of motion, see figure 1B). Changes in precision exert a substantial impact on the acuity of perception, and both humans and monkeys incorporate this impact into their decision-making (Gardelle and Summerfield, 2011; Qamar et al., 2013). Although several investigations have explored how precision impacts multisensory integration (Ernst and Banks, 2002) and perceptual learning (Adini et al., 2004; Munoz and Blumstein, 2012; Rohe and Noppeney, 2015), substantial disagreement remains as to whether and how sensory precision is reflected in subjective confidence and metacognition.

**Figure 1.**
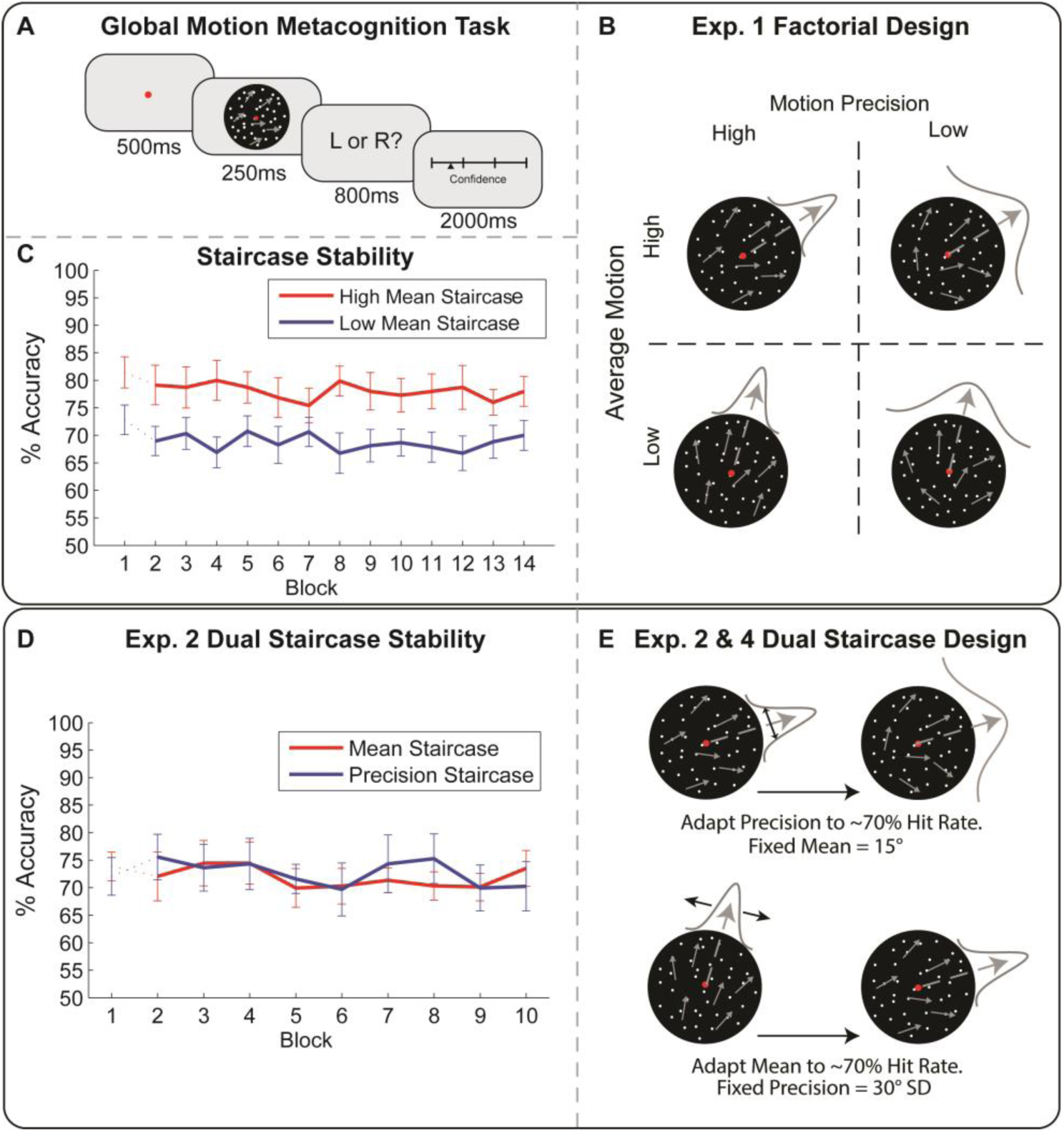
Overview of task and experimental design. **A)** We designed a global motion task to orthogonally manipulate signal mean and precision. **B)** In experiment 1, mean and precision were adjusted in a fully factorial design, to assess their impact on both perceptual and metacognitive performance. **C)** To manipulate sensory mean, we used two adaptive staircases converging on either 71% or 85% accuracy in the low vs high mean condition. Precision was manipulated by increasing the standard deviation of motion vectors for each staircase condition (i.e., by adding either 15- or 25-degrees SD to overall motion range). This adaptive thresholding procedure further stabilized detection performance across trials. Dotted lines indicate that the first block was dropped to establish staircase stabilization (high and low precision trials collapsed for illustration). **D & E)** In experiments 2 & 4, we implemented a ‘dual staircase’ design, adjusting either stimulus precision at a fixed mean or stimulus mean at a fixed precision, resulting in a low-mean, high-precision condition (mean staircase, red) and a high-mean, low-precision condition (precision staircase, blue). Experiment 3 (not shown) involved an identical task, but with two fixed levels of precision (mean-adapted to 71% hit rate). See *Methods* for more details.

For example, one early investigation found that confidence is insensitive to reductions in sensory precision, leading to over-confident orientation discrimination (Zylberberg et al., 2014). In the domain of motion perception, under-confidence in response to reduced precision has also been demonstrated (Allen et al., 2016; Spence et al., 2016; Boldt et al., 2017), and substantial inter-individual differences in the impact of sensory precision on confidence have been reported (de Gardelle and Mamassian, 2015). Collectively these studies suggest that individuals display idiosyncratic precision-related confidence biases (PRCBs); however, extant limitations preclude understanding the exact genesis and direction of these biases.

A crucial limitation is that sensory precision is typically confounded with other experimental factors, such as signal intensity and/or task difficulty, both of which can potentially impact confidence. One explanation for this influence is found in computational models of metacognition, which suggest that the brain monitors the variability of neural populations encoding a perceptual input to represent decision confidence (Meyniel et al., 2015; Spence et al., 2016; Fleming and Daw, 2017). As the intensity and precision of a visual stimulus both underpin perceptual accuracy and are likely to influence neuronal variability (Pouget et al., 2000; Knill and Pouget, 2004), we set out to disambiguate how these sources of perceptual information contribute to metacognitive confidence. To do so, we implemented a global dot-motion task specifically designed to disentangle the contributions of these factors to confidence while controlling for task difficulty (see Fig. 1B).

As previous experiments suggest that PRCBs may be driven in part by changes in physiological arousal (Allen et al., 2016; Hauser et al., 2017a), we hypothesized that precision-specific changes in confidence would correlate with changes in pupil dilation, an index noradrenergic arousal when luminance and task difficulty are controlled. In the brain, metacognitive ability (i.e., the correspondence between confidence and accuracy) is linked to individual differences in the anatomy and microstructure of the prefrontal cortex (Fleming et al., 2010; McCurdy et al., 2013; Allen et al., 2017b). However, the specific influence of sensory precision on confidence has been linked to superior parietal and ventral striatal activations (Bang and Fleming, 2018). We thus used whole-brain regression modelling to identify cortical areas whose gray-matter volume correlated with PRCBs. To address these questions, we conducted four inter-related psychophysical experiments together with physiological (Experiment 3) and neuroanatomical modelling (Experiment 4).

## Methods

### Overview

We developed a global-motion detection task for use in all four experiments (illustrated in Figure 1), in which the signal intensity and precision of the overall motion stimulus could vary independently of one another (see *Stimulus*, below). In contrast to a standard random dot kinetogram (Braddick, 1974), in which the proportion of signal (i.e., dots moving along a coherent path) and noise (i.e., randomly moving) dots is adjusted to sensory threshold, we presented motion stimuli at full coherence while independently manipulating the standard deviation and mean angle of global motion signals. In our task, motion signal intensity and precision were thus operationalized as the mean angle of all dots, and the standard deviation or range of dot-paths, respectively. Heuristically this is akin to ‘smearing’ the motion paths around an average vector; all dots have a constant average signal while the global mean and precision are independently controlled. This is a critical feature of our task, as otherwise correlations between fluctuations in mean and precision could obscure their respective influence on confidence (Zylberberg et al., 2014; Spence et al., 2016).

In all four experiments, participants performed a forced-choice discrimination and confidence-rating task after each motion presentation, while discrimination difficulty was maintained using an adaptive staircase procedure. Experiment 1 (‘factorial design’, illustrated in Fig. 1B) involved the fully orthogonal manipulation of mean and variance in a factorial design. In Experiment 2 (‘dual-staircase design’, illustrated in Fig. 1E), the impact of perceptual precision on confidence was assessed while controlling performance confounds by staircasing either stimulus mean at a fixed precision, or by adjusting precision at a fixed mean. This resulted in low-mean, high-precision (mean-staircase) and high-mean, low-precision (precision staircase) conditions while minimizing trial-wise correlations between each parameter. In Experiment 3, we adapted our paradigm for pupillometry, assessing the difficulty-independent impact of sensory precision on confidence and evoked pupil responses by adjusting stimulus mean for a high and low level of stimulus precision, similar to de Gardelle et al (2015). Experiment 4 established neuroanatomical correlates of individual differences in precision-related confidence bias; as in Experiment 2, participants completed the dual-staircase task followed by structural magnetic resonance imaging.

### Global Motion Detection and Metacognition Task

In all experiments, participants performed a speeded forced-choice global motion discrimination task. All participants were healthy young adults recruited from University College London and surrounding London community via electronic postings to a participant database and received monetary compensation for their participation. All procedures were conducted with approval from UCL research ethics committee in accordance with the Declaration of Helsinki and all participants provided written informed consent.

All experiments began with a short training task of 100-200 trials, in which a change in the colour of the fixation dot provided choice accuracy feedback on each trial, to ensure participants understood the task before continuing to the main experiment. During training trials, participants performed the motion-discrimination task without rating their confidence. Thresholds obtained for individual staircases from the training performance were then used as starting points for adaptive staircases in each experiment.

At the start of all experiments, participants were informed that the purpose of the task was to measure their perceptual and metacognitive abilities; metacognitive ability was defined in terms of their ability to introspect on each motion judgement and accurately represent their performance with confidence reports. On each trial, participants were instructed to maintain central fixation and determine the “average” or “global” direction of motion in the presented stimulus (relative to vertical) as accurately as possible. To stabilize inter-response intervals (i.e., to facilitate stable pupillometric signals, see experiment 3), participants were trained to respond with the left or right arrow keys within 800ms post-stimulus offset (Murphy et al., 2014b) in all experiments. Participants then had 2000ms to rate their confidence using a continuous visual-analogue scale, with participants moving a triangular marker along a horizontal line with the left and right arrow keys.

For reference, the confidence scale was marked by 4 vertical bisecting lines corresponding to and labelled as “No confidence”, “low”, “moderate”, and “high confidence”. For each confidence rating, participants were instructed to reflect carefully on their subjective feeling of confidence in the decision they had just made and were encouraged to use the entire confidence scale to reflect this feeling. Each trial thus consisted in a brief fixation, a motion stimulus and directional judgement, and confidence rating period lasting 2 seconds (Fig 1a). For choices faster than 100ms or slower than 800ms, a brief instruction text appeared displaying “Too fast!” or “Too slow!” and the trial was marked as missed. All missed trials were removed from any subsequent analyses, and not included in staircase updates. To prevent advance response preparation, on each trial the confidence slider’s starting point was randomly jittered by +/- 15% of the scale midpoint (Fleming et al., 2012b).

### Global Motion Stimulus

We designed a global motion stimulus to enable orthogonal manipulation of sensory mean (i.e., the average angle of dot direction), and sensory precision (the standard deviation of dot direction vectors). Across all four experiments, motion stimuli where an animated, circular display of 1,100 dots presented around a central fixation dot. All experiments were conducted in a darkened room using a 60 Hz LCD monitor and implemented in MATLAB using PsychToolbox-3 on a Windows PC; head fixation was maintained using a chin and forehead rest (Brainard, 1997; Pelli, 1997; Kleiner et al., 2007). Dots where animated in an upward-vertical direction. All stimuli were presented at 100% coherence (all dots were ‘signal’), while parametrically varying the distribution of motion vectors (i.e., sensory precision) and their orientation from the vertical axis (i.e. signal mean).

On each trial, black dots of radius 0.08 degrees visual angle (DVA) were presented for 250ms within a 15.69 DVA diameter circular array at random starting positions, with dots advancing 0.06 DVA per frame. Dots that moved beyond the stimulus aperture were replaced by dots at the opposite edge to maintain constant dot density. To prevent fixation on the local motion directions, all dots had a randomized limited lifetime of maximum of up to 14 frames, or 93% of total stimulus duration. On each trial the motion signal was thus calculated using the formula:

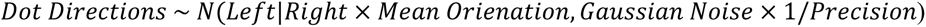

where signal mean or precision varied parametrically according to the stimulus staircase or a pseudo-randomized condition (see below for details specific to each experiment). In all experiments, equal numbers of left and right directions were presented, pseudo-randomized within each block.

In each experiment, block-wise accuracy was examined to ensure staircase stability, with the first block of trials being dropped to allow staircase stabilisation. To identify subjects whose performance failed to stabilize, we examined boxplots of mean accuracy and perceptual threshold separately for each condition, excluding participants with extremely poor performance (i.e., 1.5 times below the lower interquartile range). Further, all stimulus traces were inspected for each subject and condition to ensure adequate staircase stabilization. See individual study details, below, for study-specific exclusions.

### Experiment 1

Twenty-nine healthy volunteers (17 Female, mean age = 23 years, range = 18-40) participated in Experiment 1, in which our aim was to assess whether subjective confidence reports accurately reflect the impact of orthogonal changes of both sensory mean and precision on objective accuracy. Experiment 1 consisted of 784 trials of a global motion discrimination task in which sensory mean and precision were manipulated independently of one another in a factorial design (factors: mean, precision; levels: high/low). The task was divided into 14 blocks of 56 trials each, with the order of stimulus condition pseudo-randomized within each block.

To manipulate sensory mean, we implemented two staircases in which the average angle of motion was adjusted across trials using a staircase converging on either 71% (i.e., 2-down 1-up) or 85% accuracy (4-down 1-up). Thus, each staircase adaptively adjusted the mean signal, resulting in a lower (71% staircase) or higher mean signal (85% staircase), while maintaining stable performance within each condition. To ensure orthogonal manipulation of signal mean and precision, responses to low precision trials were not included in the staircase. Instead, on each low precision trial, orientation was generated using the signal mean from the previous low precision trial of the same (low or high) mean staircase condition. For example, on each low mean (71% accuracy staircase), low precision trial, the stimulus mean was drawn from the previous low mean, high precision trial; similarly, for each high mean (85% accuracy staircase), low precision trial the stimulus mean was drawn from the previous high mean, high precision trial. This resulted in a fully orthogonal 2 by 2 factorial design with conditions: low mean low precision (LMLP, signal μ = 3.54°, SD = 0.82°, signal precision σ = 25°), high mean low precision (HMLP, signal μ = 5.73°, SD = 2.04°, signal precision σ = 25°), low mean high precision (LMHP, signal μ = 3.57°, SD = 0.86°, signal precision σ = 15°), and high mean high precision (HMHP, signal μ = 5.73°, SD = 2.04°, signal precision σ = 15°). We note that although this resulted in a slight (0.03°) difference in average signal intensity between the LMLP and LMHP condition, this difference was not significant (p = 0.33). Boxplot inspection identified four participants for whom staircases failed to stabilize. These participants were removed from all subsequent analyses leaving a final N = 25. After removing the first block of trials, repeated measures ANOVA (factors: sensory mean, sensory precision, block) revealed good staircase stability, with no overall effect of block on detection accuracy, F(12, 24) = 0.421, p = 0.96, and no block by condition interactions (all p > 0.16).

### Experiment 2

In experiment 1, we focused on the fully orthogonal manipulation of sensory mean and precision, demonstrating their impact on perceptual sensitivity and confidence. However, factors influencing task difficulty, such as cognitive or perceptual load, have a well-known impact on confidence (Fleming et al., 2012a), we sought to replicate the precision-related confidence bias in a second task equating task difficulty while minimizing trial-wise correlation between our sensory parameters. To do so, we implemented a psychophysical ‘dual staircase’ task, in which either the mean was fixed, and the precision was adjusted using a 2-down 1-up staircase, or precision was fixed with the mean adjusted by a similar procedure. Precision- and mean-adapted trials where randomly intermixed within blocks. This enabled us to equate reaction time and perceptual sensitivity (d’), while estimating the independent effects of mean and precision on confidence.

Experiment 2 differed from the previous experiment as follows. 24 healthy, new participants (12 females, average age = 26 years, range = 18-39) participated in Experiment 2. To manipulate stimulus precision independently of task difficulty, we now employed two separate 2-up-1-down staircases with either a fixed mean (μ = 15°) and adjusted precision, or a fixed precision (σ = 30°) and adjusted mean. This resulted in a low precision, high mean condition (precision staircase, μ = 15°, σ = 48.12°) and a high precision, low mean condition (mean staircase, μ = 6.48°, σ = 30°). Critically, compared to previous investigations where both low and high precision stimuli were staircased by adapting mean signal (introducing a high trial-wise correlation of both mean and precision) (Zylberberg et al., 2014; de Gardelle and Mamassian, 2015), in Experiment 2 only mean or precision fluctuated within each condition. Participants completed 400 trials divided evenly between each staircase condition, with pseudo-randomized blocks of 40 trials each. Boxplot inspection identified four subjects in whom staircases failed to stabilize. These participants were removed from all subsequent analyses, leaving a final N = 20. After removing the first block of trials, repeated measures ANOVA (factors: block, staircase) again revealed good staircase stability: no significant main effect of block (F(8, 152) = 1.33, p = 0.23) or block by staircase condition interactions (F(8, 152) = 0.88, p = 0.54) were observed.

### Experiment 3 – Pupillometric Correlates of PRCB

In Experiment 3, we adapted our task for pupillometry to assess how task-evoked changes in arousal encode the interaction of sensory precision and confidence. Experiment 3 (N = 32, 19 females, average age = 24 years, range = 18-38) was identical to Experiment 1, with the following exceptions designed to maximize precision of our pupil measurements. To equate discrimination difficulty, which can have a profound impact on pupil dilation (Kahneman and Beatty, 1966), in two separate 2-down 1-up staircase conditions we continuously adapted mean signal with a fixed precision of either 15 or 25 standard degrees. This resulted in a low mean, low precision condition (Signal μ =3.29°, SD = 1.35°, Signal Precision σ = 25°) and a high mean, high precision condition (Signal μ = 5.96°, SD = 2.87°, Signal Precision σ = 15°). Note that this procedure introduces a trial-wise correlation between mean and precision (similar to de Gardelle and Mamassian, 2015); we subsequently control for these effects by using signal mean, accuracy, and reaction times as covariates in all our behavioural and pupillometric analyses. Boxplot inspection identified four subjects in whom staircases failed to stabilize, who were removed from all subsequent analyses, resulting in a final N = 28. For our blockwise accuracy analysis, Mauchly’s test indicated that the assumption of sphericity was violated (*p* < 0.05); therefore, we report p-values and F-statistics using the Greenhouse-Geisser correction. Repeated measures ANOVA (factors: block & precision) again demonstrated that following removal of the first block of trials, staircase stability was achieved as indicated by no main effect of block (*F*(4.27, 115.21) = 2.06, p = 0.09) or block by precision interaction (F(4.06, 109.71) = 0.72, p = 0.56).

Furthermore, as stimulus luminance is a primary driver of the pupillary response we altered the visual appearance of our stimulus to minimize frame-to-frame changes in luminance. The monitor was first calibrated with a Minolata CS-100 photometer and linearized in software, giving a mean and maximum luminance of 42.5 and 85 cd/m^2. Black dots were presented on a grey background at mean luminance. Further, following previous random-dot motion pupillometry studies (Murphy et al., 2014b; Allen et al., 2016), we implemented an isoluminant dot-masking procedure; each trial began with a dot-motion stimulus for 250ms; at motion offset a second random dot mask appeared. Participants then made their L/R motion judgement and confidence rating while centrally fixating on the dot-mask; a black confidence scale was then superimposed over the dot mask. To limit eye-movements during confidence reports, the width of the rating scale was restricted to half that of the motion stimulus. Participants completed 480 trials divided evenly between each staircase condition, with 8 randomized blocks of 60 trials each.

### Experiment 4 – Neuroanatomical Correlates of PRCB

Experiment 4 (N = 49, 28 females, average age = 24 years, range = 18-39) was identical to Experiment 2 in all aspects; new participants completed the dual-staircase version of our task, and underwent anatomical magnetic resonance imaging. Data from these subjects were previously analysed in a quantitative MRI study to examine microstructural correlates of overall metacognitive ability (Allen et al., 2017b); here we specifically investigated the precision-related bias in relation to grey matter volume. Boxplot inspection identified five participants in whom staircases failed to stabilize, who were removed from all subsequent analyses. Additionally, one subject was unable to complete the MRI acquisition resulting in a final N = 43. Repeated-measures ANOVA (factors: block, staircase) again revealed that following exclusion of block 1, no effect of block (F(6, 43) = 0.30, p = 0.94) or block by staircase interaction (F(6, 43) = 1.04, p = 0.40) was observed, indicating good staircase stability. Participants completed the metacognition task, as well as a behavioural battery probing auditory perceptual abilities, empathy, and general metacognitive ability (Allen et al., 2017a, 2017b; Carey et al., 2017). In a separate one-hour scanning experiment, participants underwent anatomical and functional brain imaging (the latter not presented here).

### MRI Acquisition

All imaging data were collected on a 3T whole body MR system (Magnetom TIM Trio, Siemens Healthcare, Erlangen, Germany) using the body coil for radio-frequency (RF) transmission and a standard 32-channel RF head coil for reception. As part of a previous investigation, we collected whole-brain quantitative multi-parameter maps (MPM), which can be used to derive traditional volumetric measures in addition to quantitative tissue maps (e.g., as markers of cortical iron and myelination). The full acquisition and preprocessing has been previously described (Allen et al., 2017a, 2017b; Carey et al., 2017), here we only recount the basic imaging details.

To this end, we acquired a whole-brain quantitative MPM protocol consisting of 3 spoiled multi-echo 3D fast low angle shot (FLASH) multi-echo gradient echo acquisitions with 800 µm isotropic resolution and 2 additional calibration sequences to correct for inhomogeneities in the RF transmit field (Lutti et al., 2010, 2012). Differential T1-weighting was achieved in two of the FLASH acquisitions by varying the flip angle (either 6° or 21°), while the third had magnetisation transfer weighting, achieved through the application of a Gaussian RF pulse 2 kHz off resonance with 4ms duration and a nominal flip angle of 220° prior to a 6° on-resonance excitation. The field of view was 256mm head-foot, 224mm anterior-posterior (AP), and 179mm right-left (RL). Gradient echoes were acquired with alternating readout gradient polarity at eight equidistant echo times ranging from 2.34 to 18.44ms in steps of 2.30ms using a readout bandwidth of 488Hz/pixel to minimise distortions. Only six echoes were acquired for the MT-weighted acquisition to maintain a repetition time (TR) of 25ms for all FLASH volumes. To accelerate the data acquisition, partially parallel imaging using the GRAPPA algorithm was employed with a speed-up factor of 2 in each phase-encoded direction (AP and RL) with forty integrated reference lines. Total acquisition time for this protocol was less than 30 mins.

The acquired data were combined to generate maps of magnetisation transfer saturation (MT) using the model described in Helms et al (2008). Following similar pre-processing protocols as used by Fleming et al. (2010) and McCurdy et al (2013), these MT maps were then segmented into grey matter, white matter, and CSF in native space. We chose to segment the MT maps because of their excellent contrast (Helms et al., 2009) and because of their greater specificity over conventional T1-weighted images (Lorio et al., 2014, 2016). The DARTEL (diffeomorphic anatomical registration through exponentiated lie algebra) algorithm (Ashburner, 2007) was used for spatial normalization to increase the accuracy of inter-subject registration, by aligning and warping the grey matter segments to an iteratively refined template space. The DARTEL template was then rigidly registered to the Montreal Neurological Institute stereotactic space, and the grey matter segments modulated such that their original tissue volumes were preserved. These grey matter segments were smoothed using an 8mm full-width at half-maximum Gaussian kernel and were manually inspected following each pre-processing step to ensure quality control. We note that in the present investigation we focused only on volumetric correlates of PRCB; see Allen et al (2017b) for results of the quantitative imaging analyses.

## Analyses – General Details

### Staircase validation analyses

We first calculated detection accuracy within each block of trials, to assess when staircase stability was reached. This was implemented using repeated measures ANOVA with factors block and condition, followed by pairwise t-tests between individual blocks to determine at which block interval performance stabilized. This analysis indicated that performance stabilized in all experiments within the first block of trials; these were subsequently removed from all further analyses. Furthermore, any missed trials were excluded. In all experiments, staircase stability was further assessed by inspecting boxplots of perceptual thresholds (median stimulus orientation or stimulus precision across trials) and accuracy separately for each condition, removing participants displaying extremely poor performance (i.e., as identified by boxplot and stimulus trace inspection).

### Signal Detection Theoretic Analyses of Perceptual and Metacognitive Performance

To quantify both type-I (perceptual) and type-II (metacognitive) performance, we applied a signal detection theoretic (SDT) approach. To do so, we first binned confidence ratings into four equal quartiles, normalizing differences in the distribution of the confidence reports. We then fit a meta-dprime (meta-d’) model to detection accuracy and binned confidence reports, to estimate the SDT parameters d’ (perceptual sensitivity), c (perceptual criterion or bias), meta-d’ (metacognitive sensitivity), and m-ratio (metacognitive efficiency, d’/m’). Heuristically, this model quantifies the sensitivity of confidence reports to objective detection performance, or P(high confidence | correct response) vs P(high confidence | incorrect response). While meta-d’ indicates the sensitivity of confidence to performance, and is akin to a “type 2” receiver-operating curve measure of metacognition, it can be confounded by differences in detection performance. However, because the model expresses meta-d’ in the same (signal detection) units as d’, the ratio (meta-d’/d’) can be calculated to quantify metacognitive efficiency, i.e., the sensitivity of confidence reports to the available perceptual signal. The meta-d’ model was thus estimated separately for all conditions using a maximum likelihood approach, implemented in the Type-II SDT toolbox (http://www.columbia.edu/~bsm2105/type2sdt/) for Matlab (Maniscalco and Lau, 2012). Furthermore, mean confidence was calculated as an estimate of type-II or metacognitive bias, i.e., the tendency for participants to report high or low confidence irrespective of performance.

### Trial-wise Prediction of Confidence

In each experiment, to estimate the impact of sensory precision on confidence independently of perceptual difficulty, we fit regression models predicting trialwise fluctuations in confidence while controlling for stimulus intensity (mean), reaction time, and decision accuracy. To do so, we constructed design matrices of trialwise regressors encoding sensory precision while controlling for sensory mean, reaction time, and accuracy (see below for specific details in each experiment). Both dependent and independent variables were normalised to comply with the assumptions of general linear models and to enable the comparison of beta-weights (i.e., effect size) between conditions.

### Statistical Analyses – Behaviour and Neuroimaging

Statistical significance was assessed using a summary statistics approach by pooling subject-level beta-weights and performing a Bonferroni-corrected one-sample t-test on each regressor of interest. All ANOVAs were conducted in JASP (Version 0.8.1.1, https://jasp-stats.org/); all t-tests and regression models in MATLAB (R2016a). Pupil data were imported using Fieldtrip (r7276) (Oostenveld et al., 2010) and analysed using MATLAB. All neuroimaging pre-processing steps and analyses were conducted using SPM12 (http://www.fil.ion.ucl.ac.uk/spm/software/spm12/).

## Experiment-Specific Analyses

### Experiment 1

Mean confidence, M-ratio, RT, d’, and perceptual thresholds (median orientation) were entered into separate repeated measures ANOVAs (Factors: Mean, Precision; Levels: High, Low). Planned pairwise t-tests were carried out comparing the difference in confidence, RT, and d’ for the low mean, high precision vs high mean, low precision conditions. Finally, the precision related-confidence bias was assessed by fitting a multiple regression model predicting trialwise confidence. The model was of the form:

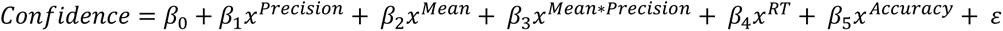

Where *x* denotes the vector of the respective independent variable across all trials, independently for each participant. This allowed us to assess the significance of the β-parameter across subjects, in a summary-statistics one-sample t-test, controlling for the influence of RT, accuracy, and stimulus mean. P-values were corrected for multiple comparisons using Bonferroni correction.

### Experiment 2

The analysis of Experiment 2 was identical to Experiment 1, with the following exceptions. Here, perceptual thresholds (i.e., the convergence point for each staircase) were calculated as either median sensory precision (precision staircase) or median sensory mean (mean staircase), as these variables where adjusted only within their respective conditions (see Fig. 1E). Paired-sample t-tests were used to assess significant group-level differences in confidence, RT, d’, and M-ratio between the two staircases. To assess the specific influence of sensory precision on confidence, we again fit a trialwise multiple regression model to the confidence data. Here the model was identical to that of experiment 1, with the exception that we did not include the interaction of sensory mean and precision as these were not manipulated in a factorial design and hence where not orthogonal to one another. Using a summary statistics approach, we then performed one-sample t-tests on each beta-parameter, using a Bonferroni correction for multiple comparisons.

### Experiment 3

As in Experiment 2, mean confidence, RT, d’, threshold (median signal orientation), and M-ratio were analysed using paired-samples t-tests comparing the high and low precision staircases. To estimate the noise-related bias, trialwise regression models predicting confidence were constructed and analysed as in Experiment 2, controlling for sensory mean, reaction time, and accuracy. All pupil pre-processing and analyses used similar procedures as in our previous investigation of how arousal impacts the precision-related bias (Allen et al., 2016). Pupil data were first imported and pre-processed in MATLAB using the Fieldtrip package (Oostenveld et al., 2010). Data for the high and low precision condition were epoched according to motion onset, from −1000ms baseline to 4000ms (rating offset). Blinks were automatically detected as any sample in which amplitude dropped below 600 arbitrary dilation units and linearly interpolated. Blinks at the beginning or end of a trial were interpolated according to the first reliable sample, and any trial with more than 25% blinks was removed from future analyses. Unreliable data due to partial eye-closures (eye fluttering or ‘mini-blinks’) were detected as signal fluctuations greater than 30 AU within a 5ms window and linearly interpolated within a 200ms time window; trials with excessive (>30 ‘miniblinks’) were rejected from analysis. All trials were de-trended and low-pass filtered using a 2-way Butterworth filter at a 30Hz cut-off, and manually inspected for remaining artefacts. These procedures resulted in an average 15.17% trial rejection rate; 8 participants were excluded from pupil analyses due to excessive rejection rates (>40%).

Pupillary time series were analysed using a trial-level general linear modelling approach. To do so, each trial was first baseline corrected for the pre-stimulus interval. Design matrices were then constructed with explanatory regressors encoding the main effects of stimulus precision, confidence, and their interaction. We also modelled stimulus mean, accuracy, and RT for each trial, to control for possible confounding effects of perceptual difficulty. Thus, at each sampled time bin, we fit a regression model of the form:

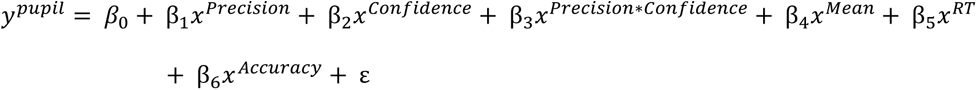

This generated beta-weight time-series for each regressor, which we then submitted to group-level permutation t-tests in a summary-statistics approach to assess when the encoding of an experimental variable (e.g., precision or confidence) significantly diverged from zero. We thus tested the consistency of the individual timeseries at the group level by conducting t-tests for the positive and negative effect of each regressor, corrected for multiple comparisons using a standard cluster-based permutation test (p < 0.05 cluster-extent correction, based on 10,000 permutations and height threshold *t* = 2.49 corresponding to a one-sided inclusion p < 0.01) (Hayasaka et al., 2004; Hunt et al., 2013; Hauser et al., 2015; Allen et al., 2016), using custom MATLAB code. This approach allowed us to assess when a particular condition or interaction of interest was significantly encoded in pupillary fluctuations, maximizing temporal sensitivity without relying on the assumptions of a deconvolution approach (Wierda et al., 2012).

### Experiment 4

All behavioural analyses were identical to those of Experiment 2. To perform voxel-based morphometry, pre-processed (segmented, normalized with modulation, and smoothed) MT-derived grey matter segments were analysed in a multiple regression design matrix to identify brain regions whose grey matter volume correlated with the precision-induced confidence bias, as derived from individual regression models predicting confidence. To control for any possible between-subject differences in perceptual performance, we also included covariates encoding the average difference in signal threshold between the precision and mean-staircases, the conditional difference in perceptual sensitivity, overall perceptual sensitivity, as well as nuisance covariates encoding age and gender.

Proportional scaling was used to account for variability in total intracranial volume across participants. To exclude clusters outside the brain and limit the search volume to voxels reasonably likely to contain grey matter, a binary grey matter mask (> 0.4) was generated using the DARTEL template. We then evaluated positive and negative correlations of the PRCB using one-sample t-tests at the group level. As in our behavioural analyses, PRCB was calculated using our trial-wise regression approach, yielding beta-weights for each subject indicating the sensitivity of confidence to sensory precision while controlling for other task-related factors such as signal mean and reaction time. Further, to explore individual differences in the influence of sensory precision on perceptual performance, we further analysed the correlation of individual differences in mean-thresholds (i.e., the amount of extra signal intensity needed to equate performance for noisy stimuli), between the high and low precision conditions with cortical volume. Statistical significance was assessed using a whole-brain, peak-corrected family-wise error (FWE) rate p < 0.05. Anatomical labels for significant brain-behaviour correlations were generated using the SPM Neuromorphometrics toolbox.

## Results

### Results Summary

Across four experiments we found that low sensory precision (high uncertainty) led to a reduction in confidence above and beyond the effect of precision on perceptual accuracy or response speed, indicating a precision-induced conservative confidence bias. In Experiment 3, we further found that this bias is related to changes in baseline arousal; exploratory computational modelling of this effect using the hierarchical drift diffusion model (HDDM) revealed that baseline arousal and sensory precision exerted independent influences on evidence accumulation. Finally, in Experiment 4 we found that the volume of the superior parietal lobe correlated with the magnitude of the conservative PRCB.

### Experiment 1

In Experiment 1 (n = 29), we manipulated sensory precision and signal intensity (mean orientation) orthogonally, allowing each to independently impact perceptual performance so that we could observe their individual and additive effects on confidence. To establish how sensory mean and precision impact perceptual performance, we analysed their effect on decision speed and sensitivity. This revealed that both stimulus mean (main effect mean, F(1, 24) = 125.00, p < 0.001, partial η2 = 0.84), precision (main effect, F(1,24) = 158.85, p < 0.001, partial η2 = 0.87), and their interaction (F(1,24) = 22.07, p < 0.001, partial η2 = 0.48) substantially impacted perceptual sensitivity (d’). This pattern of results was also reflected in median reaction times with a main effect of mean (F(1, 24) = 27.84, p < 0.001, partial η2 = 0.54); precision (main effect, F(1,24) = 29.775, p < 0.001, n2 = 0.56) and a significant interaction (F(1,24) = 22.52, p < 0.001, partial η2 = 0.48). Thus, degradations in both stimulus intensity and precision super-additively increased task difficulty (Fig. 2A), as expected by signal detection theory (Green and Swets, 1966).

**Figure 2.**
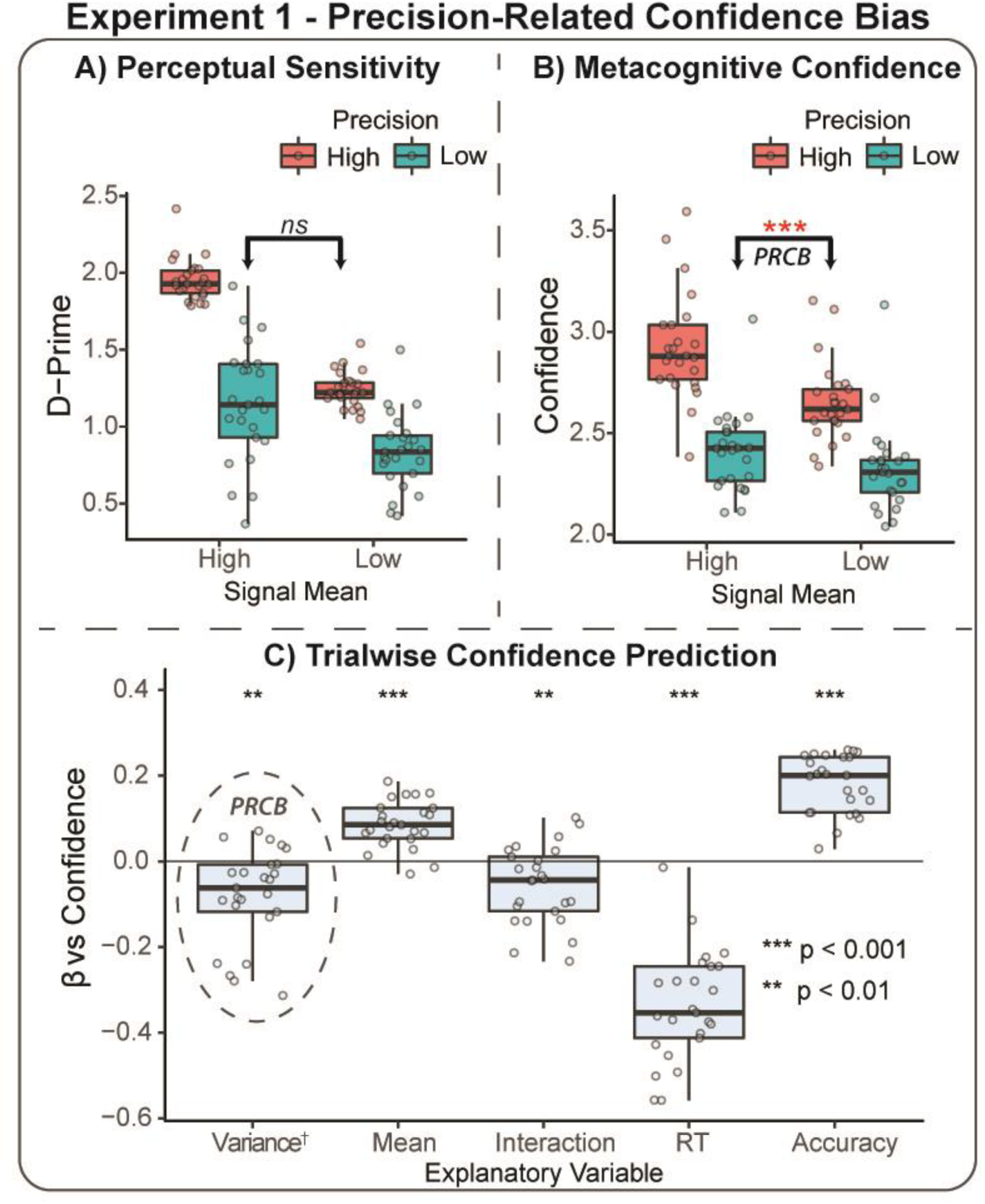
Experiment 1 Results – Precision Related Confidence Bias. **A)** Boxplots depicting motion detection sensitivity (d-prime) for each stimulus condition. Reductions in average motion signal and sensory precision both degrade perceptual performance. **B)** This pattern is amplified in subjective confidence reports, which show a strong super-additive interaction between sensory precision and mean. However, when performance is equated (middle two conditions), confidence is significantly reduced for the low-precision stimuli. This result confirms that *precision acts to bias confidence reports* in a manner that differs from the impact of precision on accuracy, i.e., a conservative precision-related confidence bias (PRCB). All main effects and interactions significant, p < 0.001. **C)** Trial-wise multiple regression predicting confidence fluctuations demonstrates that the PRCB is independent of general task performance (i.e., Reaction Time & Accuracy) and stimulus mean. Data points depict individual subject values; boxplots depict median value and interquartile range. Significance determined using summary-statistics one-sample t-tests (Bonferroni corrected). †For ease of interpretation, precision is plotted as variance (1/precision), rendering more negative beta-values as a more conservative PRCB.

We next evaluated the degree to which this effect was reflected in subjective confidence. Repeated-measures ANOVA revealed that self-reported confidence was highly sensitive to fluctuations in both mean and precision: we observed significant main effects of mean (F(1, 24) = 39.08, p < 0.001, partial η2 = 0.62, Fig. 2B), precision (main effect precision, F(1,24) = 59.39, p < 0.001, partial η2 = 0.71), and their interaction (F(1,24) = 33.25, p < 0.001, partial η2 = 0.58). It is of note that whereas sensory mean and precision exerted similarly large effects on accuracy (partial η2 = .84, .87), precision exerted a nearly 20% greater effect on confidence than did mean (partial η2 = 0.62, 0.71), indicating a substantial precision-related confidence bias.

These results indicate that confidence is sensitive to the super-additive impact of both perceptual mean and precision, providing evidence for a precision-related confidence bias. However, it is unclear if these effects on confidence are mediated by similar changes in perceptual sensitivity/difficulty or reflect a true bias in confidence. To address this question, we performed post-hoc paired t-tests on confidence for the “high mean low precision” and “low mean high precision” conditions, which did not significantly differ in terms of reaction time (t(24) = 1.85, p = 0.08) or perceptual sensitivity (t(24) = −1.24, p = 0.28). These analyses demonstrate that in spite of equivalent performance on these conditions, confidence was substantially reduced for the lower precision/higher uncertainty condition (t(24) = −4.42, p < 0.001, Cohen’s d = −0.88, Fig. 2B). Thus, even when performance is equated, participants show a substantial conservative bias (reduced confidence) for low precision stimuli. This result confirms that perceptual precision acts to bias confidence reports in a manner that differs from the impact of precision on accuracy.

Next, to determine the specificity of this bias, we performed a multiple regression analysis within each subject, predicting trial-wise confidence fluctuations from sensory precision while controlling for mean, reaction time, the precision by mean interaction, and choice accuracy. One-sample t-tests (two-sided, Bonferroni corrected for 5 comparisons) over the beta-weights for each normalized predictor showed that, independently of these other factors, lower sensory precision degraded subjective confidence (t(24) = −3.66, average beta = −0.08, p = 0.001, Fig. 2C). Additionally, all other parameters were significant; sensory mean increased confidence (t = 7.89, p < 0.001), the precision by mean interaction reduced confidence (t = −3.11, p = 0.004), lower accuracy reduced confidence (t = −13.17, p < 0.001), and higher RT correlated with reduced confidence (t = 13.35, p < 0.001).

Finally, we assessed whether sensory precision and mean also influenced metacognitive efficiency (M-Ratio). This analysis found no differences in metacognitive efficiency for stimulus mean, precision, nor their interaction (all p > 0.35), indicating that perceptual mean and precision exert independent effects on confidence (type-II bias), rather than metacognitive efficiency itself.

### Experiment 2

Experiment 2 (n = 24) was designed to test whether the precision-related bias remains when perceptual performance is equated between high and low precision conditions (Fig. 1E). Pairwise t-tests confirmed that perceptual sensitivity and median reaction times were successfully equated between the two staircases (Fig. 1D, all p > 0.13). Nevertheless, confidence was significantly reduced for the low-precision stimuli (t(19) = 2.68, p = 0.015, d = 0.60, Fig. 3A), again demonstrating that precision biases confidence independently task difficulty. No differences in metacognitive efficiency were observed (p = 0.414). These results replicate a precision-induced confidence bias as independent from both perceptual and metacognitive sensitivity.

**Figure 3.**
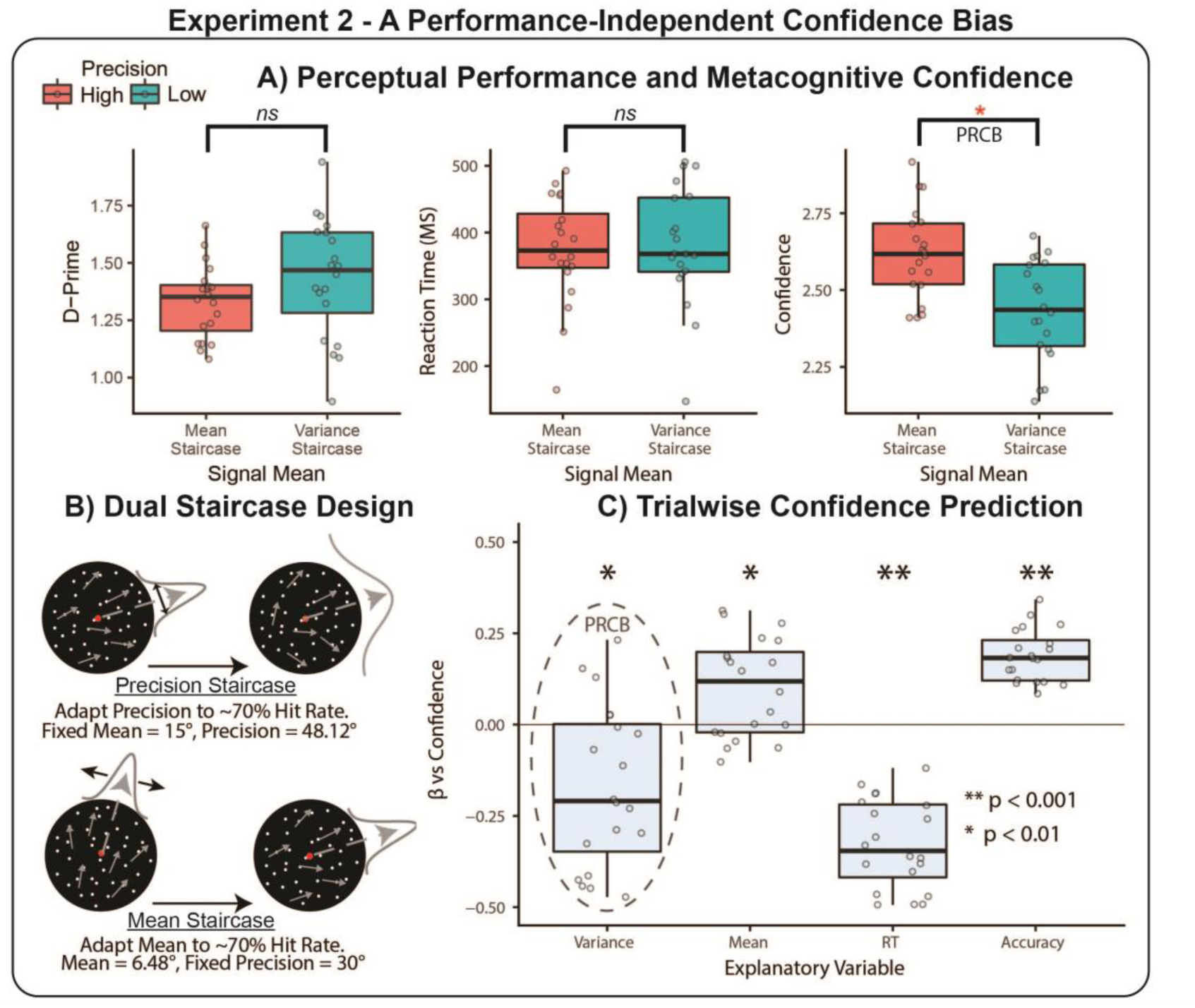
Experiment 2 replication and extension of the PRCB. **A)** The dual-staircase design successfully equates perceptual difficulty; neither motion sensitivity (d-prime) or reaction time differ significantly between staircases. Nevertheless, confidence is reduced for the lower-precision stimuli, replicating the PRCB. Data points depict individual subject values; boxplots depict median value and interquartile range. **B)** Schematic demonstrating the dual-staircase design; two independent staircases adaptively adjust either sensory precision or mean to equate performance; the precision staircase results in a lower overall sensory noise. **C)** Trialwise regression again demonstrates the independence of PRCB from task difficulty; low sensory precision reduces confidence while accounting for mean, RT, and accuracy. Significance determine using summary statistic one-sample t-tests for each regressor, Bonferroni adjusted. †For ease of interpretation, precision is plotted as variance (1/precision), rendering negative beta values a more conservative PRCB.

As in Experiment 1, to capture the independence of a precision-related confidence bias from accuracy, RT, and sensory mean, we again fit regression models predicting trial-wise confidence. This analysis reproduced the independence of the precision-related confidence bias (t(19) = −3.45, p = 0.0027, mean beta = −0.17), as well as demonstrating significant effects of mean (t(19) = 3.34, p = 0.0035), reaction time (t(19) = −12.02, p < 0.001), and accuracy (t(19) = 11.80, p < 0.001). This experiment therefore replicated the difficulty-independent, precision-related confidence bias using a novel staircase procedure, while minimizing across-trial correlation between mean and precision.

### Experiment 3

In Experiment 3 (n = 32), we optimized our global motion task for pupillometry (see *Methods*), to assess relationships between confidence, sensory precision, and their correlation with pupil dilation. To do so, we staircased high and low levels of sensory precision by adjusting mean stimulus intensity separately for each condition. This enabled us to account for differences in perceptual difficulty (a strong driver of pupillary response) in the same manner as a previous experiment (de Gardelle and Mamassian, 2015); however this approach introduces a strong trialwise correlation between mean and precision which we sought to account for in our analyses. In our behavioural analysis, we first assessed whether our procedure equated performance. Indeed, pairwise t-tests did not reveal any significant differences in mean RT (*t*(27) = 1.96, p = 0.057) or d’ (t(27) = −0.91, p = 0.37). In contrast to our previous experiments in which trial-wise mean and precision were de-correlated, here we observed no significant overall effect of stimulus precision on confidence (t(27) = 0.91, p = 0.37), replicating De Gardelle et al (2015).

As this effect is confounded by the staircase-induced correlation between sensory mean and precision, we re-analysed the confidence data in a one-way repeated measures ANCOVA (Factor: Precision), while controlling for the difference in median signal orientation between each condition. This revealed a significant main effect of precision on confidence (F(1, 26) = 8.09, p = 0.009, partial η2 = 0.19) and a precision by orientation difference interaction (F(1,26) = 7.74, p = 0.01, partial η2 = 0.19). Thus, when accounting for the staircase-induced mean difference, the precision-related bias is again apparent. This result was further confirmed by our trial-wise regression analysis predicting confidence: low precision again significantly reduced confidence (t(27) = −2.95, p = 0.007) when accounting for stimulus mean, RT, and accuracy. These results underscore the importance of accounting for both sensory mean and precision when assessing their influence on perceptual confidence.

Previous investigations have suggested that arousal contributes to perceptual confidence (Lempert et al., 2015; Hauser et al., 2017a), and that the manipulation of arousal reduces precision-related confidence biases (Allen et al., 2016). We therefore examined the correlation of sensory precision, confidence, and their interaction with pupil dilation while controlling for sensory mean, RT, and decision accuracy using a cluster-corrected general linear modelling approach (see *Analysis* for further details). This analysis revealed significant arousal-related correlates of precision, confidence, and their interaction. High sensory precision correlated significantly with increased pupil dilation between 1.45 and 2.61 seconds post-stimulus onset, maximum *t*(22) = 23.69, *p*Cluster = 0.003. Replicating previous results (Lempert et al., 2015; Allen et al., 2016), increased confidence was related to reduced dilation from 1.43s – 1.72s post-stimulus onset, minimum *t*(22) = −11.02, *p*Cluster = 0.048). Crucially, we observed a significant positive interaction of sensory precision and confidence encoded in pupil dilation during the pre-stimulus baseline (1s – 0.68s pre-trial onset, maximum *t*(22) = 4.82, *p*Cluster = 0.037); post-stimulus effects did not survive correction for multiple comparisons. These results show that the trialwise interaction of sensory precision and confidence is related to pre-stimulus pupil fluctuations, suggesting that baseline arousal is a potential source of the precision-induced bias.

**Figure 4.**
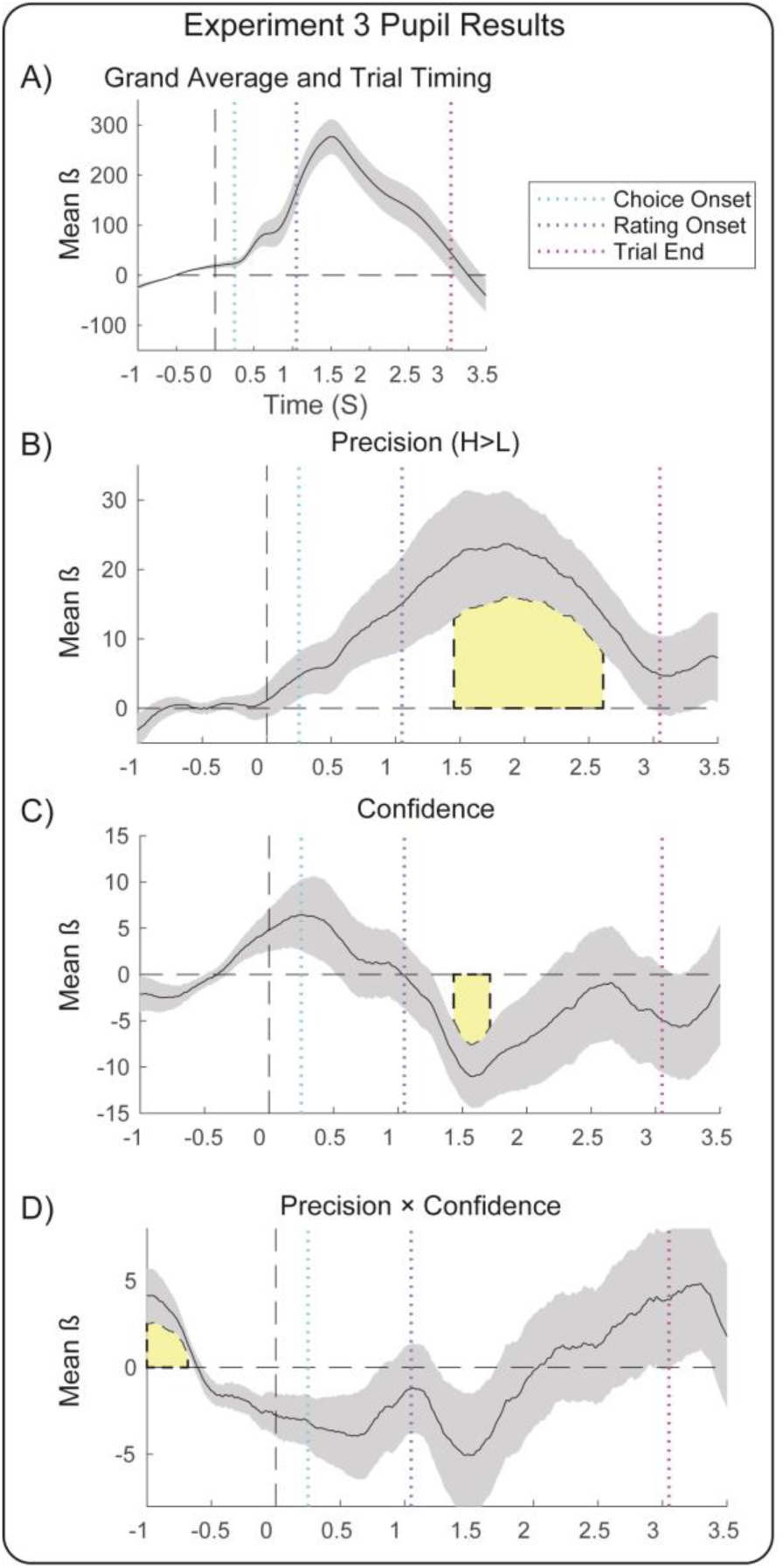
Changes in pupil dilation encode precision, confidence and the precision-induced bias. Linear modelling of evoked pupil dilation reveals significant encoding of stimulus precision, confidence, and their interaction in pupil dilation. **A)** Model intercept (i.e., the grand average) depicting trial timing relative to overall pupillary response. **B)** High sensory precision increases pupil dilation during the confidence-rating interval. **C)** Replicating previous findings (Lempert et al., 2015; Allen et al., 2016), confidence is negatively associated with dilation during the post-decision interval. **D)** A positive confidence by precision interaction is encoded in the pre-stimulus baseline, indicating that greater pre-trial dilation relates to a stronger influence of precision on confidence. Summary-statistic beta-weight timeseries from trial-level regression vs pupil response; all effects controlled for trialwise effects of sensory mean, reaction time, and choice accuracy. Significance (shaded yellow patch) determine by cluster-based permutation t-test, pCluster < 0.05 (see *Methods* for more details).

### Hierarchical Drift Diffusion Modelling of Perceptual Decision Making

We further explored the relationship between baseline arousal, sensory precision, and perceptual decision-making in our task using a drift diffusion model (DDM) to assess the impact of these variables on decision-related mechanisms. DDMs model the relationship between response accuracy and reaction time as a drifting accumulator that undergoes a biased random walk towards one or other choice boundary (e.g., left vs rightward motion) (Fig. 5A). The model thus captures a putative evidence accumulation process underpinning perceptual decision-making (Ratcliff and McKoon, 2007; Kiani and Shadlen, 2009). Notably, evidence accumulation signals recorded from the parietal cortex of macaques have been shown to relate closely to perceptual confidence (Kiani and Shadlen, 2009; Kiani et al., 2014).

**Figure 5.**
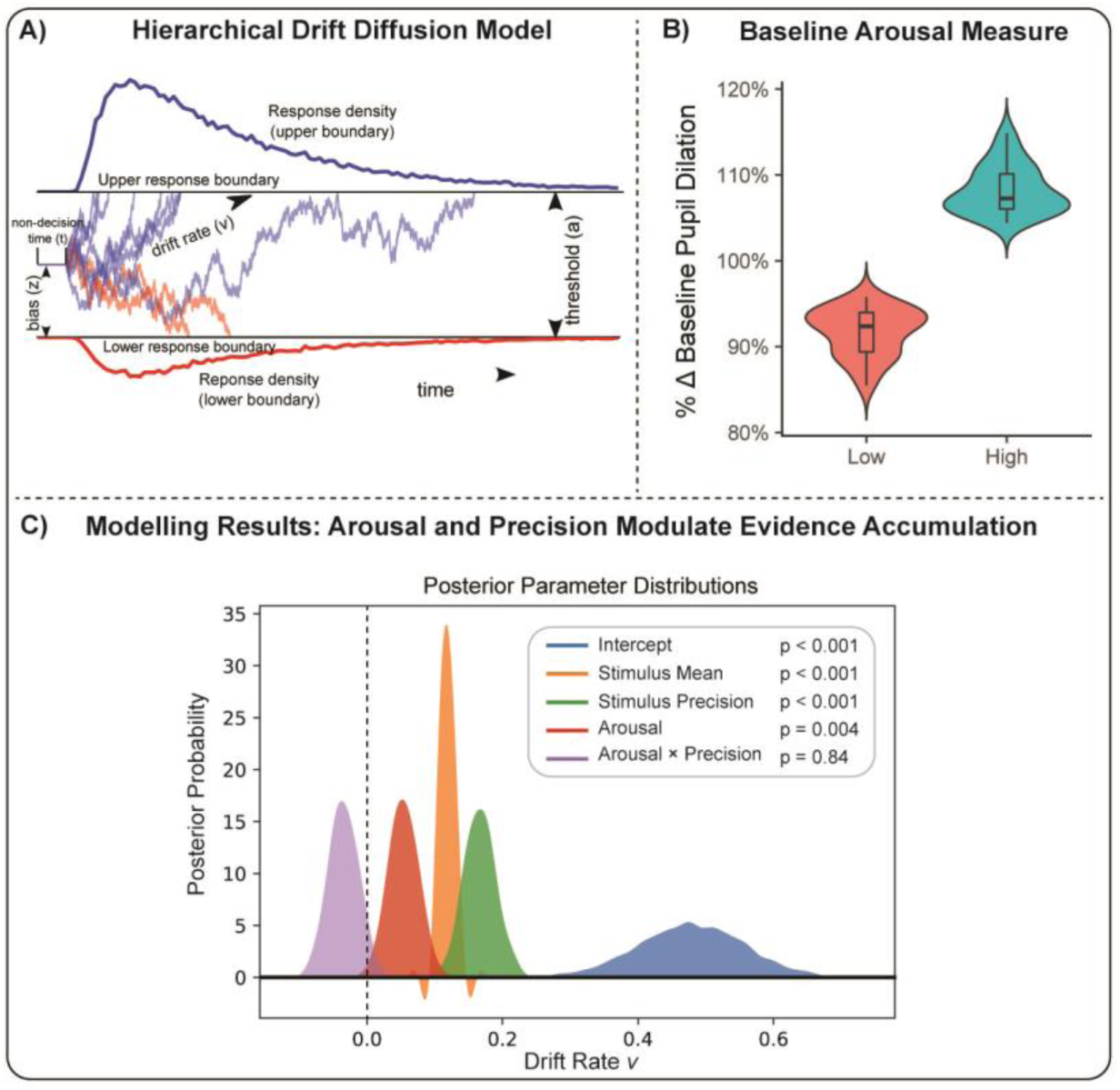
Computational Modelling of Arousal and Perceptual Decision Making. A) To better understand the observed relationship between baseline arousal, sensory precision, and confidence we fit computational models (HDDM) describing the influence of these variables on evidence accumulation. B) Baseline arousal was modelled by binning pre-trial pupil dilation into low versus high arousal conditions. We then compared models where precision, arousal, and their interaction influence the drift rate (*v*) of evidence accumulation, decision threshold (*a*), or non-decision time (*t*). C) We found that the best model was one in which stimulus mean, precision, and arousal all positively boosted evidence accumulation. However, no interaction effect of arousal and precision was observed.

To do so, we used a hierarchical drift diffusion model (HDDM) to assess the impact of pre-trial arousal and sensory precision on different model parameters. Following previous investigations (de Gee et al., 2017), ‘low vs ’high’ baseline arousal was parameterized by binning mean pre-trial dilation (−1000 to −500ms pre-trial interval) into the lowest and highest 40% of trials (Fig. 5B), separately for each participant. We then modelled the effects of sensory precision and arousal on decision making using an HDDM. The HDDM uses a Monte-Carlo Markov Chain (MCMC) method to fit a Bayesian mixed-effects implementation of the DDM, pooling over fixed and random effects to provide more robust parameter estimates. Using standard settings in the HDDM toolbox, we specified models with the following free parameters: the drift rate *v* denotes the speed of evidence accumulation, decision threshold *a* models the information threshold required to make a decision, and the non-decision time *t* captures the decision-independent processing time

Convergent evidence suggests that confidence depends on a second-order ‘read-out’ of evidence accumulation (Moran et al., 2015; Fleming and Daw, 2017); a confidence bias may thus emerge if exogenous (i.e., stimulus precision) or endogenous (i.e., arousal) factors influence underlying evidence accumulation. Because both arousal and sensory precision can be expected to increase the gain or excitability of motion-sensitive neuronal populations (Cheadle et al., 2014; Mather et al., 2015; Vinck et al., 2015), which is in turn related to evidence accumulation itself (Aston-Jones and Cohen, 2005; Eldar et al., 2013; Hauser et al., 2016), we hypothesized that the interaction of these factors covary with evidence accumulation (*v*) and that this super-additive effect may explain the over-representation of precision in confidence. To test this hypothesis, we compared models in which precision, baseline arousal, and their interaction could covary with either *a* (M2), *t* (M3), or *v* (M4). Crucially, because we controlled for task difficulty by adjusting signal strength (mean orientation), we used a regression-based approach that further allowed *v* to covary with signal strength across every trial. This enabled us to estimate the independent effect of sensory precision on different decision-related computations, and follows from previous studies demonstrating that stimulus strength directly modulates evidence accumulation (Hauser et al., 2017b). Further, we also included a ‘baseline model’ in which only *v* could covary with signal strength (M1). Models were compared using deviance information criterion (DIC); the individual posterior parameters of the best-fitting (i.e., with the best trade off of complexity to accuracy) model where then assessed for statistical significance. All models were fit with 500 burn-in samples and 10,000 total samples; inspection of posterior model parameters revealed good model convergence for all models using these parameters. See Table 1 for model comparison results.

**Table 1.**
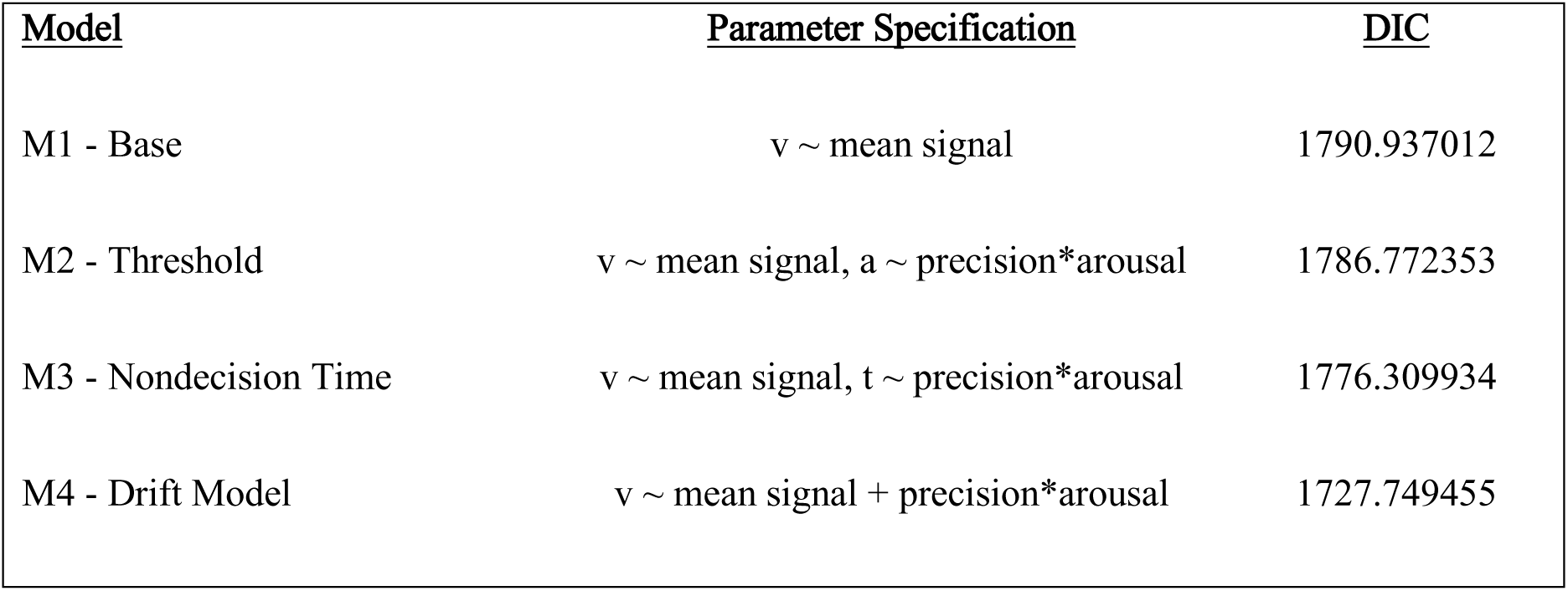
HDDM Model Comparison Results. Five models (M1-M5) where fit in which either decision threshold (*a)*, non-decision time (*t*), or drift rate (*v*) covaried with precision, arousal, and their interaction. In all models, drift rate was allowed to covary with stimulus mean signal. The best fitting model (M4) was determined by Deviance Information Criterion (DIC).

| Table 1 |  |  |
| --- | --- | --- |
| *HDDM Model Comparison Results* |  |  |
| <u>Model</u> | <u>Parameter Specification</u> | <u>DIC</u> |
| M1 - Base | $v \sim \text{mean signal}$ | 1790.937012 |
| M2 - Threshold | $v \sim \text{mean signal}, a \sim \text{precision} * \text{arousal}$ | 1786.772353 |
| M3 - Nondecision Time | $v \sim \text{mean signal}, t \sim \text{precision} * \text{arousal}$ | 1776.309934 |
| M4 - Drift Model | $v \sim \text{mean signal} + \text{precision} * \text{arousal}$ | 1727.749455 |

We found that the best-fitting model was one in which stimulus precision, baseline arousal, and their interaction covaried with *v* (see Table 1). Inspection of the posterior parameters of this model (Fig. 5C) revealed that both sensory precision (p < 0.001) and arousal (p = 0.004) significantly increased evidence accumulation (drift rate). Replicating previous investigations, stimulus mean also increased drift rate (Hauser et al., 2017b); however, in contrast to our hypothesis the arousal by precision interaction was not significant (p > 0.05). These results support previous investigations that link arousal to boosts in perceptual evidence accumulation (Lufityanto et al., 2016), and indicate that sensory precision and arousal both independently contribute to evidence accumulation but do not interact.

### Experiment 4

In Experiment 4 we applied our dual-staircase task, illustrated in Figure 1D, to identify a neuroanatomical basis for precision-related confidence bias. Previous experiments have demonstrated that inter-individual differences in overall metacognitive ability are related to prefrontal brain volume and microstructure (Fleming et al., 2010; McCurdy et al., 2013; Allen et al., 2017b); however, a recent functional MRI study suggests that the superior parietal cortex and ventral striatum may support the impact of sensory precision on confidence (Bang and Fleming, 2018). We thus sought to identify neuroanatomical correlates of inter-individual differences in precision-specific confidence bias, using an identical (dual-staircase) task and regression analyses as in Experiment 2.

Behavioural analysis showed that confidence was again significantly reduced for the lower-precision stimuli (t(43) = −5.40, p < 0.001) despite similar perceptual performance. Our trialwise regression analysis further replicated the independence of this effect from sensory mean, RT, and accuracy, with a significant negative impact of reduced precision on confidence (mean beta = −0.27, t(43) = −4.58, p < 0.001). We then used the beta-weight encoding the specific influence of precision on confidence as an index of the confidence bias in each participant, in a whole-brain multiple regression voxel-based morphometry analysis.

These analyses revealed a significant positive correlation between inter-individual differences in the precision-related bias and the volume of the left superior parietal lobe (centred at MNI coordinates XYZ = [−20, −66, 33], peak t =5.66, z = 4.76, cluster extent *k* = 435 voxels, pFWE = 0.023). This indicates that participants with lower grey matter volume in the left superior parietal lobe were more sensitive to perceptual precision, displaying a stronger reduction of confidence in response to reduced sensory precision. See Figure 6 for illustration of these results.

**Figure 6.**
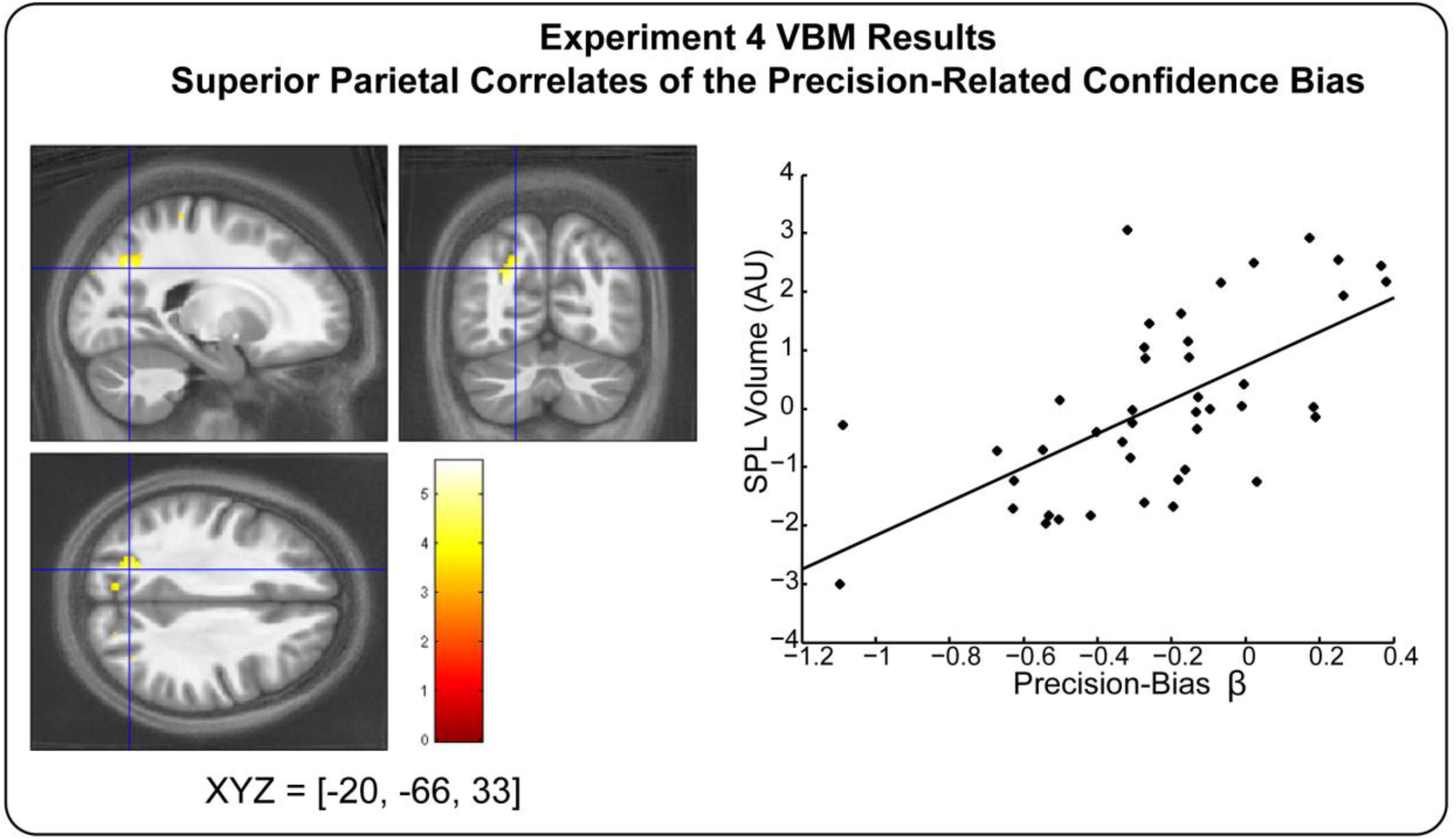
Neuroanatomical correlates of precision-related confidence bias and perceptual performance. SPL grey matter volume correlates with individual differences in precision-bias. Replicating experiment two, the precision-related noise bias was estimated using trialwise confidence regression in a dual staircase design. This enabled us to relate the precision-bias to inter-individual variability in grey matter volume. The precision bias correlates positively with the volume of the left superior parietal lobe, indicating that a more conservative bias (more negative beta-value) relates to reduced SPL volume Yellow blobs depict SPMs overlaid on a grand average MT map. Colour bar depicts t-value at each voxel. All effects significant at pFWE < 0.05 peak level correction. For illustration purposes, right panels depict bivariate correlations across individuals after adjusting for nuisance covariates.

## Discussion

Perception is an inherently noisy, imprecise process. As such, accurate perceptual choice depends upon both the sensitivity of an observer to sensory signals and on the uncertainty inherent in perceiving such signals. Similarly, accurate metacognition in the presence of uncertainty requires the ability to flexibly adjust confidence to reflect the influence of precision on our perception. Across four experiments we found that sensory precision exerts an outsized influence on subjective confidence, and that this biasing effect is at least partially independent of choice difficulty, processing speed, and signal intensity (Exp. 1-4). Further, we find that inter-trial and inter-individual variability in this effect exposes novel neurobiological correlates as measured by pupillary markers of autonomic arousal (Exp. 3) and the neuroanatomy of right superior parietal cortex (SPC, Exp. 4). These results suggest that both endogenous and exogenous factors contribute to the influence of precision on perceptual metacognition.

Across all experiments, decreases in sensory precision exerted a conservative bias on subjective confidence reports. In general, the impact of precision on confidence was substantial; observed effect sizes ranged from medium to large demonstrating that participants systematically overweight sensory precision in their estimates of average perceptual motion; further, this bias was independent of confounding factors such as sensory intensity (mean orientation), reaction time, or accuracy. These findings corroborate and extend recent behavioural reports of an overly conservative confidence bias in response to decreased perceptual precision (Allen et al., 2016; Spence et al., 2016; Boldt et al., 2017). However, other studies reported precision-related over-confidence (Zylberberg et al., 2014) or inter-individually heterogeneous biases with no overall sign (de Gardelle and Mamassian, 2015). Of these, the former was based upon a static line orientation discrimination task and the latter did not independently manipulate motion precision and mean signal. In contrast, conservative biases have typically been reported when using motion stimuli (e.g., Spence et al., 2016) and/or when controlling for signal mean (Boldt et al., 2017). Indeed, in Experiment 3 we used a similar staircasing procedure to de Gardelle and Mamassian, which introduces a high correlation of mean and precision, and found no apparent precision-bias unless the impact of sensory mean was accounted for. As such it appears both that precision-related biases may be at least partially idiosyncratic to the particular domain of perception (e.g., motion vs static orientation) and further underscores the importance of controlling for the additive effect of perceptual intensity and precision when estimating such biases.

To illuminate the neurophysiological underpinnings of this bias, we conducted pupillometric analyses together with computational modelling. In the absence of changes in luminance or choice difficulty, pupillary fluctuations are assumed to be linked to the noradrenergic output of the locus coeruleus and overall physiological arousal (Murphy et al., 2014a; Joshi et al., 2016). Here, we carefully controlled for these potential confounders, replicating recent reports of an overall negative correlation between subjective confidence and pupil dilation during the post-decision interval (Lempert et al., 2015; Allen et al., 2016). We additionally discovered a significant confidence by precision interaction, such that the trial-by-trial influence of precision on confidence correlated with changes in pre-stimulus (baseline) pupillary dilation. Our computational modelling of these results further indicated that arousal and sensory precision both independently contribute to the rate of evidence accumulation.

Previous investigations have found that baseline fluctuations in cortical alpha frequency, a marker of endogenous attention and arousal, are linked to confidence biases (Sherman et al., 2016; Samaha et al., 2017), and that baseline arousal itself increases evidence accumulation (Lufityanto et al., 2016; de Gee et al., 2017). As the computation of decision evidence is itself closely linked to confidence (Kiani and Shadlen, 2009; Fetsch et al., 2014), these results suggest that metacognition likely integrates multiple sources of environmental and endogenous noise when translating decision evidence into subjective confidence. Our results thus corroborate and extend the emerging linkage of arousal, evidence accumulation, and biases in both perception and metacognition. Although there is evidence that the biasing effect of arousal on confidence is causal in nature (Allen et al., 2016; Hauser et al., 2017a), it must be noted that in the present design we can only conclude that arousal, precision, and confidence are correlated; as we do not directly manipulate arousal here, it could simply be that the observed pupillometric effects are downstream causes of changes in confidence, rather than effects.

Experiment 4 also uncovered a novel neuroanatomical basis for inter-individual differences in confidence bias in the right SPC. In Rhesus monkeys, firing rates in lateral intraparietal area (LIP), part of the SPC, are linked to confidence-related evidence accumulation (Kiani and Shadlen, 2009), and in humans, functional MRI has recently revealed that sensory precision increases brain responses in this area independent of changes in overall confidence (Bang and Fleming, 2018). Our finding extends these results by suggesting a general role for SPC in a mapping between perceptual precision and subjective confidence. Inter-individual biases in the precision-related bias thus expose idiosyncratic patterns of cortical variability in areas strongly linked to the representation of sensory evidence; this finding may enable more specific cortical phenotyping in psychiatric populations exhibiting systematic confidence biases (Hauser et al., 2017b; Rouault et al., 2018).

What mechanism underlies the precision-related bias? Although most early models of confidence emphasized a role for serial, feed-forward processing growing evidence of confidence-accuracy dissociations (Fleming et al., 2015; Allen et al., 2016) cast doubt on models which specify that the computation of confidence is wholly subsumed by the first-order decision variables governing choice accuracy itself. Instead our results may be better explained by either a parallel processing architecture and/or a top-down predictive processing scheme (see e.g., Fleming and Daw, 2017).

As an example of the former, it is possible that the visual system has evolved to attribute extra salience to fluctuations in perceptual precision. In this case, confidence may arise both from a serial architecture in which prefrontal neurons monitor the variability of parietal evidence accumulation to provide an initial estimate of confidence (Kiani and Shadlen, 2009; Maniscalco and Lau, 2016), and a parallel pathway specifically signalling perceptual salience. This is a potentially compelling explanation of our results: evoked changes in arousal are directly linked to enhanced perceptual salience (Hochstein and Ahissar, 2002) and are also known to change the gain or non-linearity of cortical responses (Simoncini et al., 2012; Eldar et al., 2013). Thus, changes in baseline arousal may simultaneously alter perceptual salience and increase the sensitivity of confidence-related brain areas to changes in perceptual precision.

Alternatively, predictive-processing accounts of metacognition posit that confidence depends upon the mismatch between an expected level of perceptual precision and actual incoming sensory precision (i.e., a confidence prediction error) (Fetsch et al., 2015; Allen et al., 2016; Allen and Friston, 2016; Fleming and Daw, 2017). As changes in baseline arousal are linked to increased levels of neuronal noise and poorer task performance (Aston-Jones and Cohen, 2005; Mather et al., 2015), such changes may alter top-down estimates of expected precision and hence titrate the influence of sensory precision on confidence. Disentangling these (speculative) suggestions will require tasks that manipulate both expected and sensory precision in conjunction with causal manipulations of arousal; here we can only demonstrate that changes in sensory precision evoke a performance-independent bias, supporting these and other non-serial metacognitive architectures.

## Conclusions

Our study provides strong confirmation of a performance-independent confidence bias in response to fluctuations in perceptual precision. This bias is found above and beyond the impact of precision on task performance, suggesting that parallel and/or top-down architectures may provide a better model of confidence than standard feed-forward models. Further, this bias is linked to fluctuations in baseline arousal and individual differences in cortical neuroanatomy. As many psychiatric illnesses involve confidence biases or metacognitive deficits (Bliksted et al., 2017; Hauser et al., 2017b; Rouault et al., 2018), a PRCB may be a useful computational marker for disordered self-monitoring in diseases such as psychosis, autism, and OCD.

## Acknowledgements

The authors thank Francesca Fardo and Martina F. Callaghan for helpful discussions and input on the manuscript, MRI sequence acquisition, design, and analysis. This work was supported by Wellcome Trust grant 100227 (MA, GR) and an ERC Starting Grant to DSS (310829). The Wellcome Centre for Human Neuroimaging is supported by core funding from Wellcome (203147/Z/16/Z). MA is supported by a Lundbeckfonden Fellowship (R272-2017-4345). RJD holds a Wellcome Trust Senior Investigator Award (098362/Z/12/Z). TUH is supported by a Wellcome Sir Henry Dale Fellowship (211155/Z/18/Z), a grant from the Jacobs Foundation, and a 2018 NARSAD Young Investigator grant (27023) from the Brain & Behavior Research Foundation. A Wellcome Trust Cambridge-UCL Mental Health and Neurosciences Network grant (095844/Z/11/Z) supported RJD and TUH. SMF is supported by a Sir Henry Dale Fellowship jointly funded by the Wellcome Trust and Royal Society (206648/Z/17/Z). The UCL-Max Planck Centre is a joint initiative supported by UCL and the Max Planck Society.

